# Population genetics and phylogeographic history of the insular lizard *Podarcis lilfordi* (Gunther, 1874) from the Balearic Islands based on genome-wide polymorphic data

**DOI:** 10.1101/2024.03.06.583662

**Authors:** Katherin Otalora, Joan Lluís Riera, Giacomo Tavecchia, Andreu Rotger, José Manuel Igual, Jean-Remi Paul Trotta, Laura Baldo

## Abstract

Islands provide a great system to explore the processes that maintain genetic diversity and promote local adaptation. We explored the genomic diversity of the Balearic lizard *Podarcis lilfordi*, an endemic species characterized by numerous small insular populations with large phenotypic diversity. Using the newly available genome for this species, we characterized more than 300,000 SNPs, merging Genotype by Sequencing (GBS) data with previously published Restriction-site associated DNA sequencing (RADSeq) data, providing a dataset of 16 island populations (191 individuals) across the range of species distribution (Menorca, Mallorca, and Cabrera). Results indicate that each islet hosts a well-differentiated population (Fst=0.247±0.09), with no recent immigration/translocation events. Contrary to expectations, most populations harbor a considerable genetic diversity (mean nucleotide diversity, Pi=0.144±0.021), characterized by overall low inbreeding values (Fis<0.1). While the genetic diversity significantly decreased with decreasing islet surface, maintenance of substantial genetic diversity even in tiny islets suggests variable selection or other mechanisms that buffer genetic drift. Maximum-likelihood tree based on concatenated SNP data confirmed the existence of the two major independent lineages of Menorca and Mallorca/Cabrera. Multiple lines of evidence, including admixture and root testing, robustly placed the origin of the species in the Mallorca Island, rather than in Menorca. Outlier analysis mainly retrieved a strong signature of genome differentiation between the two major archipelagos, especially in the sexual chromosome Z. A set of proteins were target of multiple outliers and primarily associated with binding and catalytic activity, providing interesting candidates for future selection studies. This study provides the framework to explore crucial aspects of the genetic basis of phenotypic divergence and insular adaptation.

## 1. Introduction

Island populations represent biological scenarios of greatest evolutionary interest due to their geographic isolation, which accelerates the process of organismal diversification (Warren et al., 2015; Whittaker et al., 2017), and their restricted scale of study, that allow close monitoring (Drake et al., 2002; Losos and Ricklefs, 2009). They also typically host a high rate of endemism, making them a global conservation priority (Manes et al., 2021; Sivaperuman et al., 2008).The Balearic Islands, encompassing the Pityusic (Ibiza and Formentera) and Gimnesic (Mallorca, Menorca, Cabrera and associated islets) islands, comprise a large number of islands that originated following a complex geological and climatic history related to the Mediterranean sea level variations during the Pleistocene (0.2-Mya) (Cuerda, 1989; Goy et al., 1997). Among the several endemic species, they host the two species of Balearic lizard (i.e., *Podarcis lilfordi* from the Gimnesic islands and *P. pityusensis* from the Pityusic islands). They are among the most extensively studied vertebrates of the archipelago, due to their remarkable phenotypic diversity (Pérez-Cembranos et al., 2020), high population densities and reduction of anti-predatory behavior associated to the “island syndrome” (Cooper and Pérez-Mellado, 2012; Hawlena et al., 2009; Novosolov et al., 2013, Rotger et al. 2023).

*P. lilfordi* (Günther, A., 1874) presently occurs on 43 off-the-coast islets of Mallorca and Menorca, as well as in the Cabrera archipelago. Current distribution is the result of a vicariance process during the Mediterranean sea desiccation and subsequent reflooding (5.33 Mya) (Krijgsman et al., 1999; Terrasa et al., 2009). The disappearance of *P. lilfordi* from the main islands of Menorca and Mallorca dates back more than 2000 years, following the introduction of predators by humans (Alcover, 2000). The species is now confined on islets that provide an effective refuge for to date populations. Within these islets, the species evolved in a context of limited resources and absence of major predators and competitors, greatly diversifying in morphology, pigmentation (Pérez-Cembranos et al., 2020; Rotger et al., 2021), behavior and life history traits (Salvador, A., 2009; Rotger et al., 2023). Major forces driving this diversification can be ascribed to the high intrapopulation competition (density-dependent factors) (Grant and Benton, 2000; Le Galliard et al., 2010; Massot et al., 1992), as well as intense genetic drift resulting from recurrent bottlenecks, primarily affecting the smallest islets (Bassitta et al., 2021; Charlesworth et al., 2003; Rotger et al., 2021; Terrasa et al., 2009). This context of rapid phenotypic differentiation among insular populations makes *P. lilfordi* an interesting vertebrate model for ecological and evolutionary studies (Camargo et al., 2010), particularly those addressing the genomic basis of local adaptation and persistence of small populations (Bassitta et al., 2021; Yang et al., 2022).

Genetic analyses of this species have been long limited to mtDNA, few nuclear genes (Brown et al., 2008; Terrasa et al., 2009) and microsatellites for few populations (Bloor et al., 2011; Rotger et al., 2021). A recent study based on single nucleotide polymorphisms (SNPs) data by Restriction Site Associated DNA Sequencing (RADSeq) (Bassitta et al., 2021) has expanded on previous knowledge on genetic diversity and phylogeographic pattern of this species, confirming the existence of two well-discriminated genetic lineages separating populations from the major archipelagos of Menorca and Mallorca/Cabrera (Brown et al., 2008; Terrasa et al., 2009. Recently, the *P. lilfordi* genome has been sequenced (Gomez-Garrido et al., 2023) providing the framework for a comprehensive exploration of population-level genomics of this species.

Here, we used for the first time the newly available reference genome of *P. lilfordi* to map and annotate newly generated genome-wide polymorphic markers obtained by two independent sequencing methods, Genotype by Sequencing (GBS) and RADSeq. We specifically characterized SNPs of eight new populations (100 individuals) by GBS, and integrated RADSeq data of additional 10 populations (91 individuals from Bassitta *et al*. (2021), for a comprehensive analysis of 16 populations (two were common to both studies), spanning the main range of distribution of this species. Individual GBS and RADSeq datasets were independently analyzed as well as merged to obtain a representative set of common SNPs loci across all populations, which allowed for a robust comparative genomics within *P. lilfordi*.

Our main objectives were to: a) assess the level of intraspecific genetic diversity and structuring of the different populations of *P. lilfordi*; b) reconstruct the species main phylogeographic scenario of colonization of the Balearic Islands; c) identify potential signatures of genome diversification and loci underpinning the phenotypic diversity and insular adaptation of this species.

## 2. Materials and Methods

### 2.1. Sampling and DNA extractions

Tails tissues of 100 specimens of *Podarcis lilfordi* were collected between 2015 and 2016 from eight islets in the Menorca and Mallorca archipelagos (Balearic Islands) (10-22 samples per islet) (Figure 1 and Table 1). Islets encompassed a representative subsample of the current species geographic distribution and vary in several geographic and ecological aspects, including surface area, maximum altitude, vegetation cover and type, presence of human settlements and other vertebrates, as well as lizard demographic traits (Mayol et al., 2020; Pérez-Mellado et al., 2008; Rotger et al., 2021). Specimens were captured using pitfall traps placed along paths and vegetation edges, sexed according to visual examination and morphology (Rotger et al., 2016), weighted, and body size measured as snout to vent length (SVL) (see Table S1 for specimen information). Tails were preserved in 100% ethanol and kept at −80°C until processed. Permits for sampling were provided by Conselleria d’Agricultura i Medi Ambient i Territori, Govern de les Illes Balears (CEP 31/2015 to LB and CEP 6/14 to GT). Genomic DNA was extracted with DNAeasy Tissue and Blood kit (Qiagen) with RNase treatment. DNA concentration was measured with Qubit 2 fluorometer (Invitrogen, Waltham). About 1 ug of genomic DNA per sample was sent to *Centro Nacional de Analisis Genomico* (CNAG, Spain) for sequencing. DNA integrity and concentration was further measured on a fragment analyzer (Agilent Bioanalyzer) and high-quality samples with at least 10 ng/μl were selected for sequencing.

**Figure 1:**
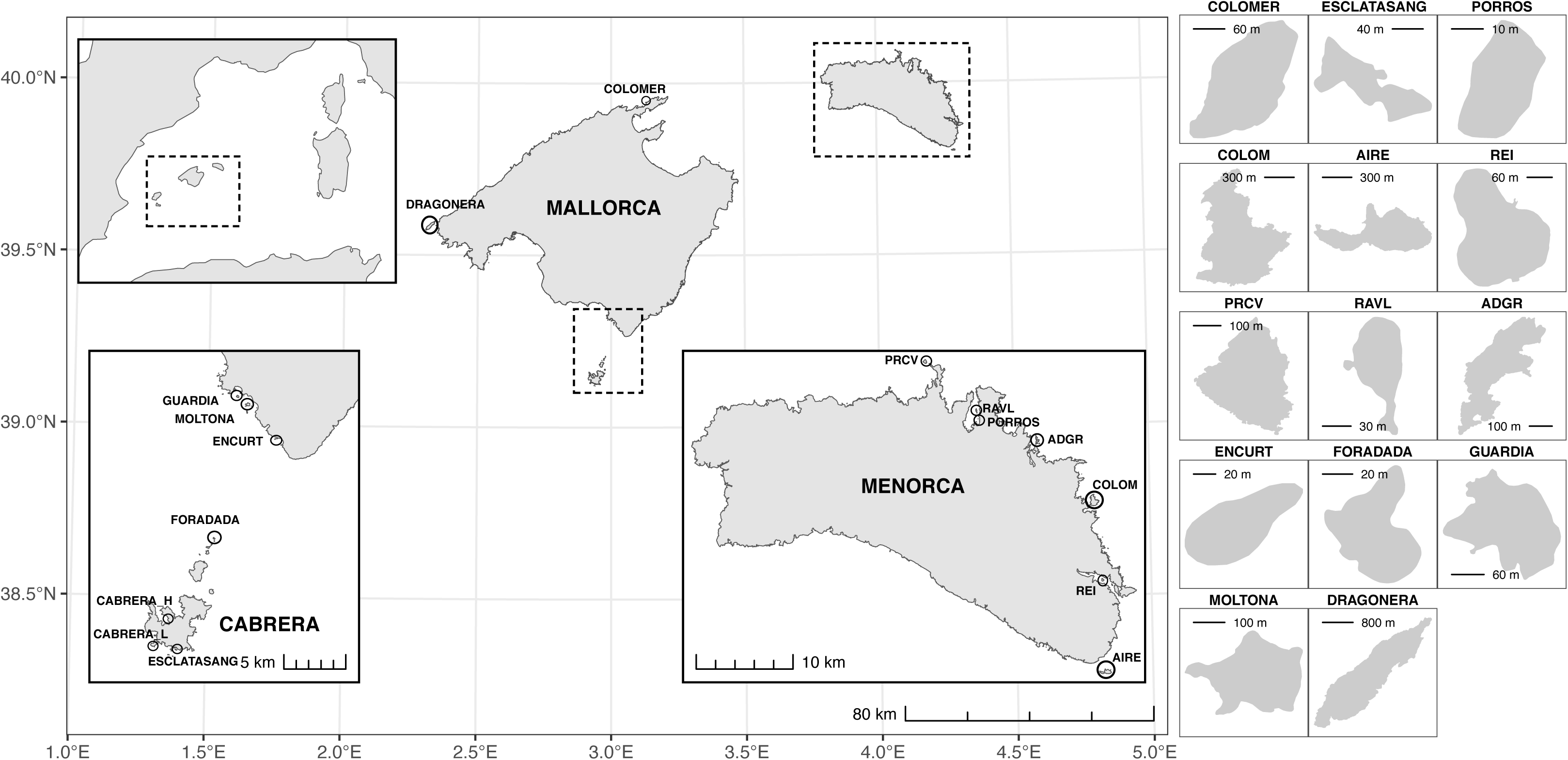
Geographic map of the 16 localities (15 islands) under study, sampled in the archipelagos of Mallorca, Cabrera, and Menorca.

**Table 1:** Sampling design and environmental variables.

| Major Archipelagos | Archipelago | Type | Official name<sup>1</sup><br>(locality) | Short name | No. of Individuals |  |  | Geographic variables<sup>1</sup> |  |  |  |  | Lizard Density<br>(ind/ha) |
| --- | --- | --- | --- | --- | --- | --- | --- | --- | --- | --- | --- | --- | --- |
|  |  |  |  |  | GBS | RADSeq<sup>2</sup> | Total | X-coord | Y-coord | Surface area (ha) | Maximum Altitude (msnm) | SDI |  |
| Menorca | Menorca | Islet | Illa Gran d'Addaia | ADGR | 10 |  | 10 | 603284,8 | 4430301,1 | 7,5 | 21,7 | 2,251 | 1062 |
| Menorca | Menorca | Islet | Illa de l'Aire | AIRE | 11 | 12 | 23 | 610443,5 | 4406455,4 | 30,5 | 16,1 | 1,946 | 4098 |
| Mallorca/Cabrera | Cabrera | Inland | Cabrera (Harbour) | CABRERA_H |  | 12 | 12 | 494459,1 | 4333606,0 | 1142,0 | 171,4 | 3,524 | 45059,0 |
| Mallorca/Cabrera | Cabrera | Inland | Cabrera (Lighthouse) | CABRERA_L |  | 8 | 8 | 493279,9 | 4331172,0 | 1142,0 | 171,4 | 3,524 | 45059,0 |
| Menorca | Menorca | Islet | Illa d'en Colom | COLOM | 10 | 10 | 20 | 609129,9 | 4424151,2 | 59,3 | 44,0 | 2,117 | 1616 |
| Mallorca/Cabrera | Mallorca | Islet | el Colomer | COLOMER |  | 4 | 4 | 511211,5 | 4421664,5 | 2,8 | 105,4 | 1,187 | 3339 |
| Mallorca/Cabrera | Mallorca | Islet | sa Dragonera | DRAGONERA |  | 4 | 4 | 441589,9 | 4381854,5 | 272,7 | 352,8 | 2,049 | 729 |
| Mallorca/Cabrera | Mallorca | Islet | Illot d'en Curt | ENCURT | 23 |  | 23 | 503125,0 | 4347776,2 | 0,4 | 3,1 | 1,194 | 1799 |
| Mallorca/Cabrera | Cabrera | Islet | Esclatasang | ESCLTS |  | 10 | 10 | 495084,6 | 4330730,1 | 0,6 | 39,2 | 1,73 | 3118 |
| Mallorca/Cabrera | Cabrera | Islet | Illot de na Foradada | FORADADA |  | 10 | 10 | 498200,0 | 4339516,6 | 0,3 | 5,7 | 1,294 | 2595 |
| Mallorca/Cabrera | Mallorca | Islet | na Guardis | GUARDIA | 15 |  | 15 | 500100,3 | 4351217,2 | 2,5 | 8,6 | 1,392 | 1784 |
| Mallorca/Cabrera | Mallorca | Islet | na Moltona | MOLTONA | 11 |  | 11 | 500926,8 | 4350573,0 | 5,7 | 7,9 | 1,395 | 2093 |
| Menorca | Menorca | Islet | Porros de Fornells | PORROS |  | 11 | 11 | 597159,9 | 4432654,7 | 0,1 | 2,2 | 1,128 | 1676 |
| Menorca | Menorca | Islet | Illa des Porros | PRCV | 10 |  | 10 | 591657,0 | 4438581,0 | 8,6 | 18,4 | 1,621 | 1100 |
| Menorca | Menorca | Islet | Illa des Revells | RAVL | 10 |  | 10 | 597032,1 | 4433305,5 | 0,4 | 8,0 | 1,518 | NA |
| Menorca | Menorca | Islet | Illa del Rei | REI |  | 10 | 10 | 610060,7 | 4415982,1 | 4,3 | 15,2 | 1,221 | 403 |

<sup>1</sup> Vector geographic information was extracted from Base Topogràfica de les Illes Balears (https://ideib.caib.es/geoserveis/rest/services/public/GOIB_BTIB_IB/MapServer/27), publicly available from Institut Cartogràfic i Geogràfic de les Illes Balears (ICGIB). Elevations were obtained from the 2-m resolution MDT02 Digital Elevation Model (DEM) publicly available from the Centro Nacional de Información Geográfica (https://centrodedescargas.cnig.es).
<sup>2</sup> from Bassitta et al. 2021

### 2.2. Sequencing

A restriction site-associated DNA library was generated by GBS method (Elshire et al., 2011). To select for the best enzyme with reproducible genomic fragments across samples (a critical criterion for GBS experiments), we first generated a pilot library (four samples) with Illumina MiSeq (150 bp), using two common restriction enzymes, Pstl and ApekI. PstI provided the best reproducibility and number of variants and was therefore chosen to generate the final library.

Genomic DNA was digested with the PstI, fragments were tagged with individual barcodes, PCR-amplified, multiplexed, and sequenced on dual lanes on an Illumina HiSeq 2500 platform (2×100 bp). Raw read sequences were demultiplexed and quality checked using FastQC v0.10.1 (Wingett and Andrews, 2018); available at https://www.bioinformatics.babraham.ac.uk/projects/fastqc/).

Additionally, raw RADseq genomic data of *P. lilfordi* generated on a previous study (Bassitta et al., 2021) were downloaded from the online repository (PRJNA645796, 91 samples from 10 populations). The GBS dataset (this study) and the RADseq dataset (Bassitta et al., 2021) shared the restriction enzyme PstI, which allowed data integration for comparative purposes (see below). Two localities of Colom and Aire were sampled and sequenced in both studies (different individuals) (Table 1 and S1).

### 2.3. SNPs calling and filtering

Raw sequence data from GBS and RADSeq were independently processed as well as merged (COMBINED dataset). Thereafter, the three generated datasets (GBS, RADSeq and COMBINED) were processed through the same pipeline.

All barcoded raw reads were mapped to the *P. lilfordi* genome (rPodLil1.1, available at https://denovo.cnag.cat/podarcis (Gomez-Garrido et al., 2023) using the Burrows-Wheeler Aligner (BWA) software (v2.1) to generate individual Binary Alignment Map (BAM) files. The genome corresponds to a single female individual sampled in the Aire population (the population was included in our dataset). Variant calling per sample was performed with Genome Analysis Toolkit HaplotypeCaller from GATK v. 3.7 (Van der Auwera GA & O’Connor BD., 2020) to produce gVCF files (minimum-mapping-quality 20). gVCF files were finally genotyped for variant calling (i.e., SNPs).

Prior to data filtering, SNP and sample statistics were explored using VCFTools v. 0.1.16 (Danecek et al., 2011; De la Cruz and Raska, 2014). SNPs were then filtered using the following settings: --max-alleles 2, --remove-indels, --min-meanDP 10, --max-meanDP 50, -- minQ 30, --maf 0.05, --max-missing 0.75 (individual datasets) or 0.90 (combined).

To retrieve common loci between GBS and RADSeq datasets, the max-missing parameter was set to 0.9, thus retaining loci shared by at least 90% of the specimens and effectively removing all loci unique to either GBS or RADSeq.

### 2.4. Population genetic diversity

We used the *populations* module in STACKS *v.2.61* (Catchen et al., 2013) to calculate diversity statistics per population, including the observed heterozygosity (Ho), expected heterozygosity (He), nucleotide diversity (Pi), inbreeding coefficient (Fis) and number of private alleles (PAs). Allelic richness (Ar) was calculated with the ‘hierfstat’ package in R (Goudet et al., 2015) using four random individuals per population, i.e. the minimum number of samples per population (minimum number of alleles, min.all=8). Significant associations between genetic diversity indexes (He, Pi, Ar, Fis) and geographic variables (islet surface area, maximum altitude, and the shoreline development index, SDI, i.e., a measure of island perimeter complexity, all log-transformed) were assessed with simple and multiple linear regression in R.

### 2.5. Population genetic differentiation and clustering

For differentiation and structural analyses, loci under Linkage Disequilibrium (LD) were removed using plink v2.00a3.3 (Purcell et al., 2007) (--indep-pairwise: 50 [’kb’] 5 0.20) (Table 2). Pairwise Fst values (Weir & Cockerham, 1984) were estimated with the function *stamppFst* from ‘StAMPP’ R package (Pembleton et al., 2013), and significant differences for each population pair were evaluated with confidence limits at 95% after 1000 bootstrap iterations. Heatmaps based on Fst were created using *pheatmap* in R (Raivo Kolde, 2019).

**Table 2:**
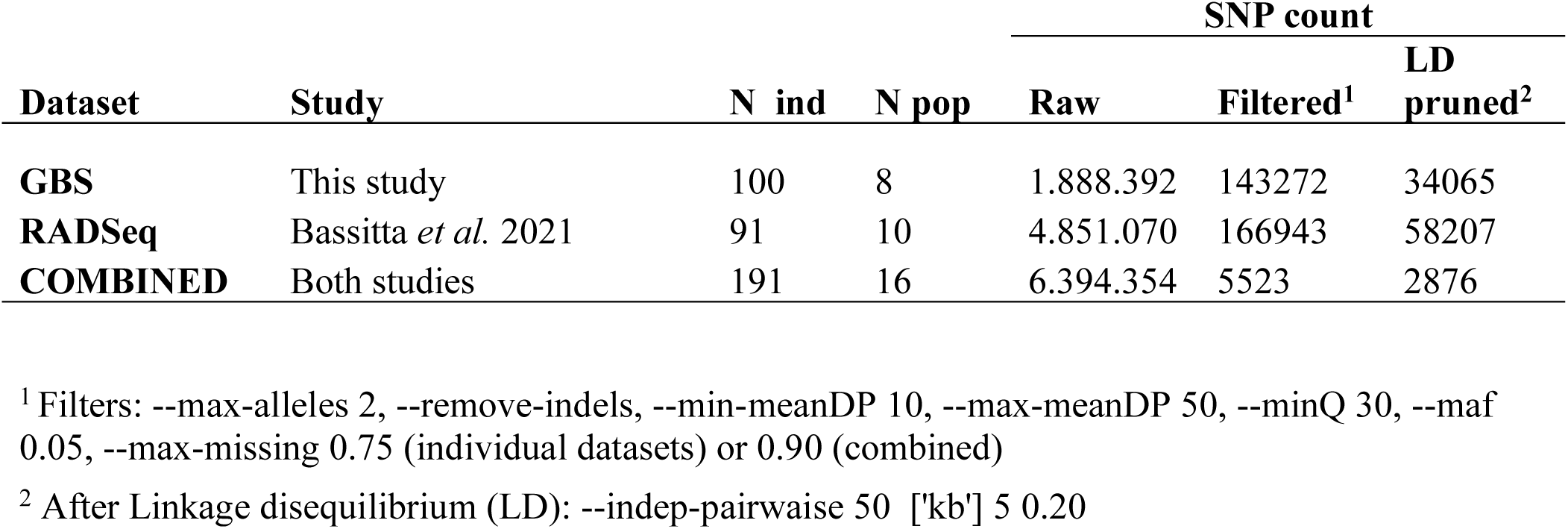
Summary of raw and filtered SNP count per dataset.

Isolation by distance (IBD) was evaluated by correlating the Fst genetic distances with linear geographic distances among populations, using a Mantel test (Mantel, 1967) with function *mantel.randtest* in the ‘ade4’ package in R (Dray and Dufour, 2007) with 1000 permutations. The test was performed for Menorca and Mallorca/Cabrera archipelago separately to control for distance effect between major archipelagos.

To reconstruct evolutionary relationships among island populations, we first used a Discriminant Analysis of Principal Components (DAPC), integrated in the ‘Adegenet’ R package (Jombart, 2008). In this multivariate statistical approach, sample genetic variance is partitioned into groups to maximize discrimination, without prior knowledge on genetic structure. The number of meaningful PCs retained was chosen by a cross-validation with the function *xvalDapc* (Jombart & Collins, 2015) with default parameters (1000 replicates).

Population structure, including contemporary and historic admixture among populations and their relationships, were further investigated using ADMIXTURE v1.3.0 (Alexander et al., 2009), setting the numbers of co-ancestry clusters (*K*) from 2 to 16 (maximum number of populations). Cross-validation was performed with the -cv option (10-fold) to retain the best value of K clusters (lowest cv error). Membership probabilities were calculated according to the retained discriminant functions and specimens showing above 90% of assignment probability to an island different from their source were considered as recently translocated, or potential mislabeled and removed (none detected).

### 2.6. Phylogeographic pattern

TreeMix v. 1.13 (Pickrell and Pritchard, 2012) was used to infer population splitting and mixture patterns. The method employs allele frequencies to build a graph-based model of the population network (as opposed to a bifurcating tree) by first building a Maximum Likelihood (ML) tree and then searching for migration events that increase the composite likelihood (Flesch et al., 2020; Pickrell and Pritchard, 2012). The program utilizes a Gaussian approximation to model genetic drift (drift parameter) along each population (Flesch et al., 2020; Pickrell and Pritchard, 2012). Using the combined data set filtered for LD, a ML tree was built with a window size (k) of 500 SNPs, evaluating from 0 to 16 migrations edges (*m*), 10 iterations per edge, using the “-noss” option to prevent overcorrection of sample size. The optimal number of significant migration edges was then inferred from the second-order rate of change in likelihood (Δ*m*) across incremental values of *m* with the *OptM* package in R (Fitak, 2021).

In addition to network analyses and as a complementary result, a Maximum Likelihood (ML) tree was built using the concatenated SNP dataset (2867 bp). Prior to concatenation, the combined VCF file was filtered using BCFtools (Li, 2011) to remove individuals with more than 10% of missing data (N=8) and candidate SNP outliers (N=18, see section 2.7). The filtered VCF file was then converted to PHYLIP format using the *vcf2phylip.py* script (Ortiz, 2019). Consensus sequences for each population were estimated with the function *consensusString* in the R package ‘Biostrings’. The two Cabrera localities were considered as a single population (Fst =0.03) and data merged. A ML tree was then built on the consensus alignment with IQ-TREE 2 v2.2.0.8 (Minh et al., 2020) using variable sites only and applying an ascertainment bias correction for SNP data (model GTR+ASC) (Lewis, 2001), with 10000 pseudo-replicates.

To identify the most ancestral population in our dataset we used IQ-TREE 2 with non-reversible substitution models (model 12.12) (Naser-Khdour et al., 2022) with 1,000 ultrafast bootstrap replicates, using both the consensus alignment as well as one random specimen per population (all sites or only variables). The program performs a bootstrap analysis to obtain several ML rooted bootstrap trees; it then computes *rootstrap* support values for each branch in the tree, as the proportion of rooted bootstrap trees that have the root on that branch. A root testing was then performed with option --root-test to compare the log-likelihoods of the trees being rooted on every branch of the ML tree. The resulting trees were visualized and edited with Figtree v1.4.4 (http://tree.bio.ed.ac.uk/software/figtree/).

### 2.7. Identification and functional annotation of SNPs outliers

To detect footprint of genomic diversification and potential selection, outlier loci were identified using two independent approaches: PCAdapt software v 4.3.5 (Luu et al., 2017) and BayeScan v 2.1 (Foll and Gaggiotti, 2008). The PCAdapt package implemented in R software detects outlier loci based on PCA by assuming that markers excessively related to population structure are candidates for potential adaptation. The number of PCs to retain was chosen after checking score plots for population structuring, setting maf=0.01 and distance to ‘mahalanobis’. The distribution of loadings (SNP contribution to each PC) was uniform, indicating no relevant LD effect. P-values were corrected for false discovery rate (FDR) using a cut-off q< 0.01 for outlier retention. BayeScan is designed to detect potential genetic loci under selection by analyzing variations in allele frequencies among specific groups with a Multinomial-Dirichlet model. The prior odd (PO) for neutrality indicates the ratio of selected:neutral sites (e.g., 1:1000) and provides a measure of uncertainty on the likelihood of the neutral model compared to the selection model (Lotterhos and Whitlock, 2014). The sensitivity of the analysis to the PO was evaluated using alternative values (1:100, 1:10000). The final MCMC chain was run for 20 short pilot runs with 5000 integrations, 50000 burn-in, thinning interval of 10, and PO set to 100. Loci were filtered for q-value < 0.01.

PCAdat and Bayescan results were finally intersected, retaining outliers with Fst > 0.8, thus providing a conservative set of candidate loci. The above analysis was performed separately for each dataset (GBS, RADseq and COMBINED data).

The candidate SNP outliers were annotated by cross-referencing the SNP position against the GFF file of the *rPodLil1.1* genome assembly (Gomez-Garrido et al., 2023) for Gene ID association. For outliers falling in protein-coding region, a Gene Ontology (GO) annotation was performed, followed by a functional enrichment analysis with gGOSt in g:Profiler [https://biit.cs.ut.ee/gprofiler/gost] on individual datasets (GBS and RADSeq). The *Podarcis muralis* genome was used as a reference to determine the functional categories (Biological processes (BP), molecular functions (MF) and cellular components (CC)) that were significantly enriched (FDR < 0.05). For the subset of outliers falling within coding regions (CDS), we annotated the codon position and assessed whether the alternative allele (ALT) translated into a synonymous or nonsynonymous substitution with respect to the reference position in the genome (REF).

### 2.8. Outlier association with environmental variables

A potential association between outliers and geographic/environmental variables (Major Archipelagos, Islet surface, Maximum altitude and SDI) was assessed through a permutational multivariate analysis of variance (PERMANOVA) with the function *adonis2* in the ‘vegan’ R package (Oksanen et al., 2022). A Euclidean distance matrix among individuals was built on allele frequencies calculated with *gl.percent.freq* in the ‘dartR’ package (Gruber et al., 2018). Previously, missing genotype values were imputed using the function ‘gl.impute’ in the dartR package. Model selection was performed by first fitting a model with all two-way interactions and all main terms, and then refitting the model after excluding non-significant interactions. Significance of model terms was assessed using marginal tests (1000 permutations). SNPs that significantly discriminated between major archipelagos were identified with logistic regression models using p<0.01 after adjusting p-values to minimize false discovery rate (FDR). A genotype matrix coded as 0/1/2 was obtained using function *extract.gt* in the ‘vcfR’ R package (Knaus and Grünwald, 2017, 2016) and used to visualize geographic clustering of individual SNP outliers by a Principal Components Analysis (PCA).

## 3. Results

We generated a total of 15 billion paired-end GBS raw reads for 100 specimens from eight distinct localities/islands (99% passing the filters, Table S2). Newly generated data and previous RADSeq data from Bassitta *et al*. (2021) were individually processed through the same pipeline and combined for an integrative analysis of all 16 populations. After genome mapping, variant calling and filtering, we obtained a total of 143272 biallelic SNPs for the GBS dataset (100 individuals, 8 localities), 166943 for the RADseq dataset (91 individuals, 10 localities) and 5523 for the COMBINED dataset (191 individuals, 16 localities, including only common loci between GBS and RADSeq datasets) (Table 2). As expected, multiallelic variants represented a minor fraction of the total polymorphic sites (2.1% for GBS and 2.5% for RADSeq) and were excluded from the study.

Both raw and filtered SNPs were widely distributed across the 18 autosomal and two sexual chromosomes (Z and W), with SNP frequencies being proportional to chromosome size (Figure 2; chromosome W representation strictly depends on the number of males in the dataset, see Table S1). Both sequencing methods covered the entire genome diversity, with chromosome representation being highly congruent among the three datasets (Figure 2).

**Figure 2:**
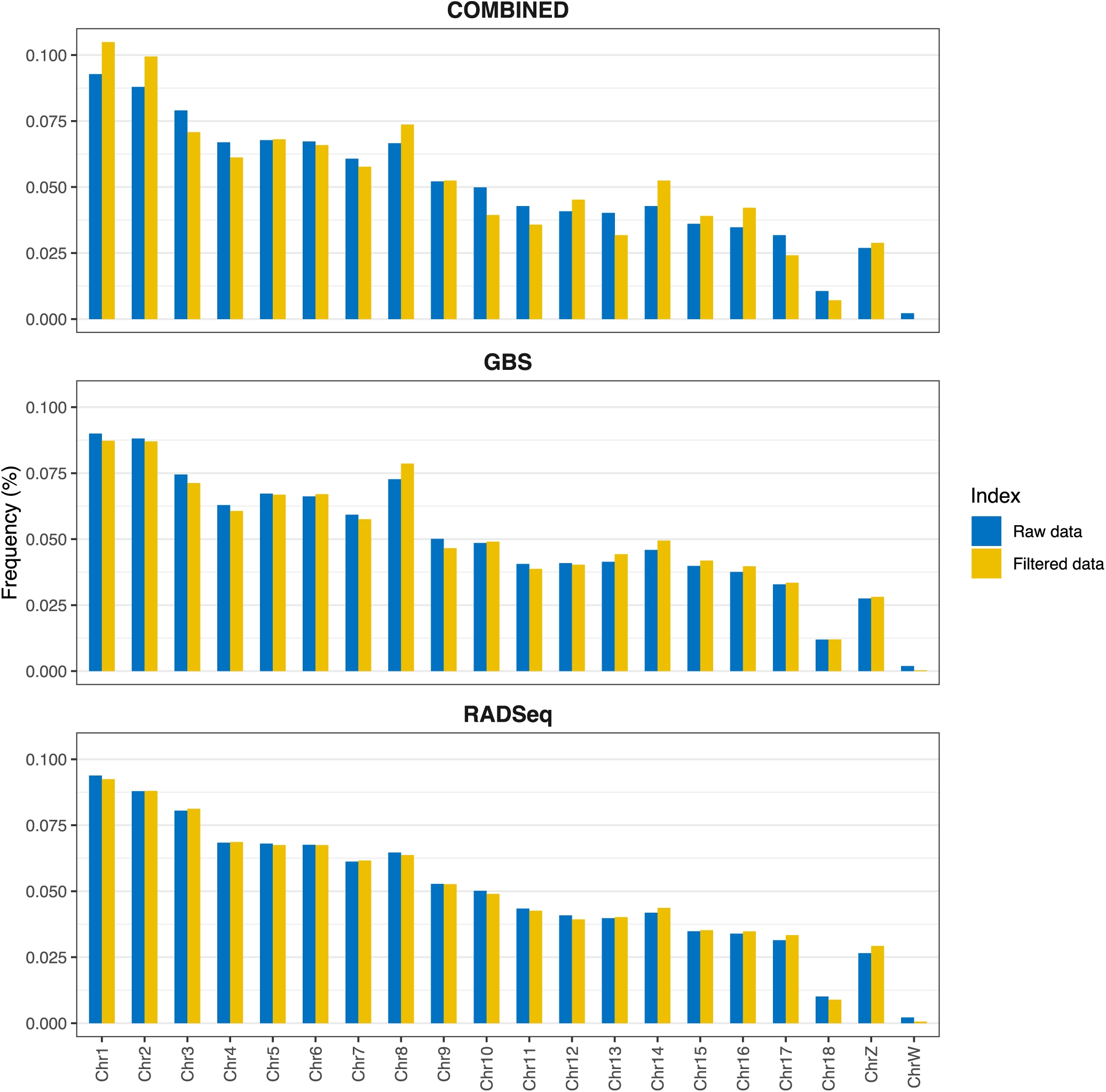
Chromosome distribution of raw and filtered SNP data for each of the three datasets (COMBINED, GBS and RADSeq).

### 3.1. Population/Islet genetic diversity

Genetic diversity based on the COMBINED dataset is summarized in Figure 3. While diversity statistics for the individual datasets (GBS and RADSeq) was slightly higher in the absolute estimates per locality (Table S3), the relative pattern found among localities was highly preserved, indicating that the reduced COMBINED dataset is overall representative of the relative genetic diversity within this species.

**Figure 3:**
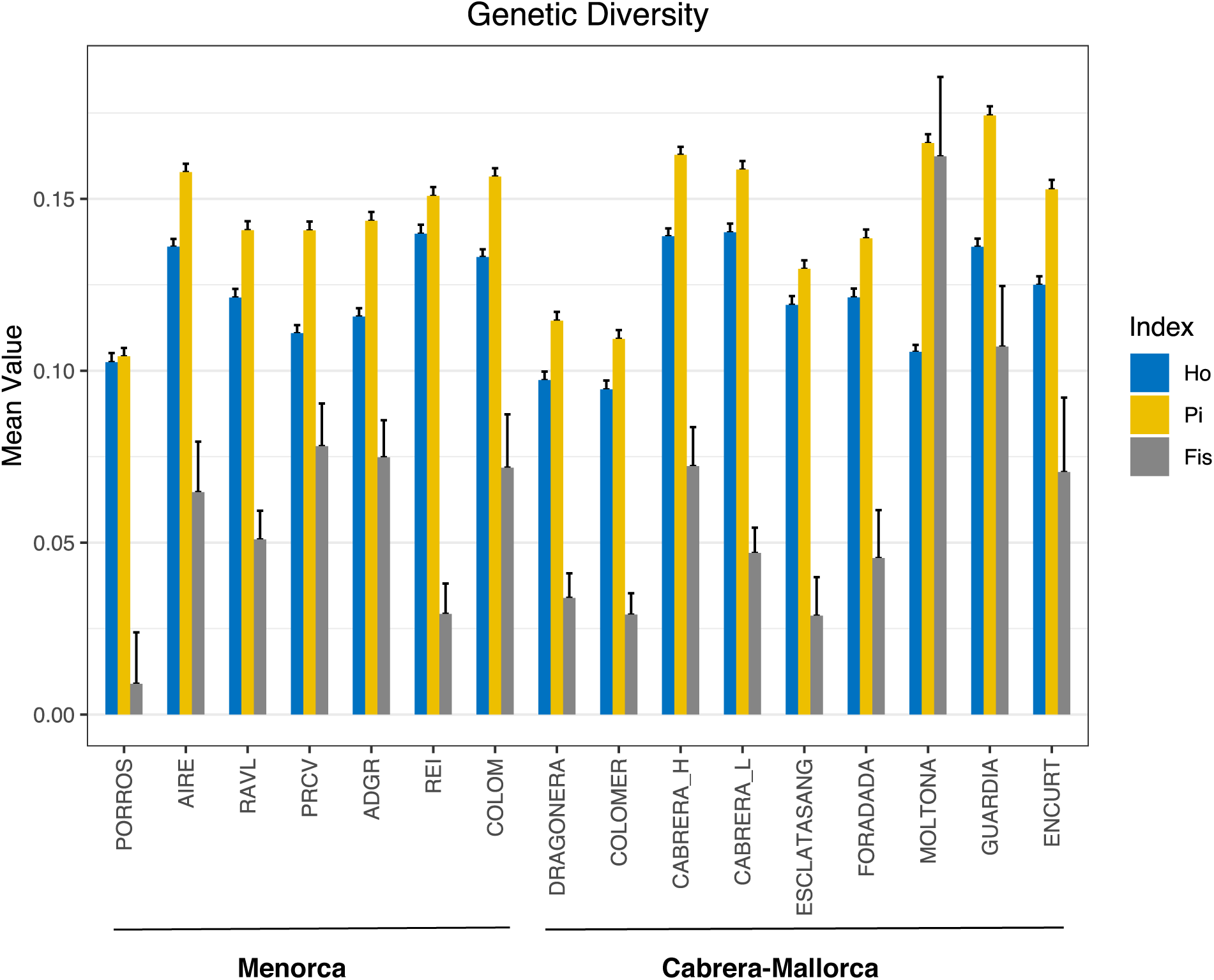
Genetic diversity estimates per population (Pi: nucleotide diversity, Ho: Observed heterozygosity, and Fis: inbreeding coefficient) with corresponding error bars, based on 5523 SNPs (COMBINED dataset).

Population nucleotide diversity (Pi) ranged between 0.104 and 0.174, with the three archipelagos showing similar ranges: 0.123-0.156 (Cabrera), 0.109-0.174 (Mallorca) and 0.104-0.158 (Menorca) (Figure 3). The observed heterozygosity (Ho) showed comparable values as Pi, varying between 0.095 and 0.140. Most diverse populations/islets per archipelago were the two Cabrera localities (Cabrera), Guardia and Moltona (Mallorca), and Aire and Colom (Menorca); the least diverse populations were the small islets of Porros (Menorca), Colomer (Mallorca) and Esclatasang (Cabrera) (Figure 3). Fis values were positive for all populations and range from 0.009 (Porros) to 0.107 (Moltona), indicating low inbreeding effect despite the small size of some islands. Individual datasets (GBS and RADSeq) based on a more extensive number of SNPs confirmed low Fis values and major trend among populations (Table S3). The Allelic richness (Ar) was also comparable across localities, ranging between 1.245 (Porros) and 1.438 (Dragonera). The number of private alleles (PA) estimated on individual datasets (Table S3) vary greatly, with highest values found in the four smallest islets of Porros, en Curt, Esclatasang and Foradada (n>1000, island area < 1.50 ha), with the small islet of en Curt presenting the largest number (n=5316) (Table S3).

Of the genetic diversity indexes, Ar, Pi and Fis showed positive coefficients with the islet surface and negative with maximum altitude in multiple regression models (Ar: p<0.001 only for Area, Pi: p=0.001 for Area and p<0.005 for Max altitude, Fis: p<0.05 for Area and Max altitude), indicating that the smallest islets typically host a more reduced genetic diversity, although no evident inbreeding depression.

### 3.2. Population genetic differentiation and structure

Pairwise Fst values (Weir & Cockerham, 1984) estimated for the combined dataset (2876 SNPs after LD pruning, Table 2) indicated that all populations were significantly differentiated (p<0.01 for all pair-wises) (Figure 4). Global levels of differentiation among islet populations were generally high, with a mean Fst *=*0.247 (ranging between 0.030 and 0.473). These values were highly comparable to those estimated on individual datasets, indicating that this reduced SNP dataset is also representative of the genetic differentiation among the studied populations.

**Figure 4:**
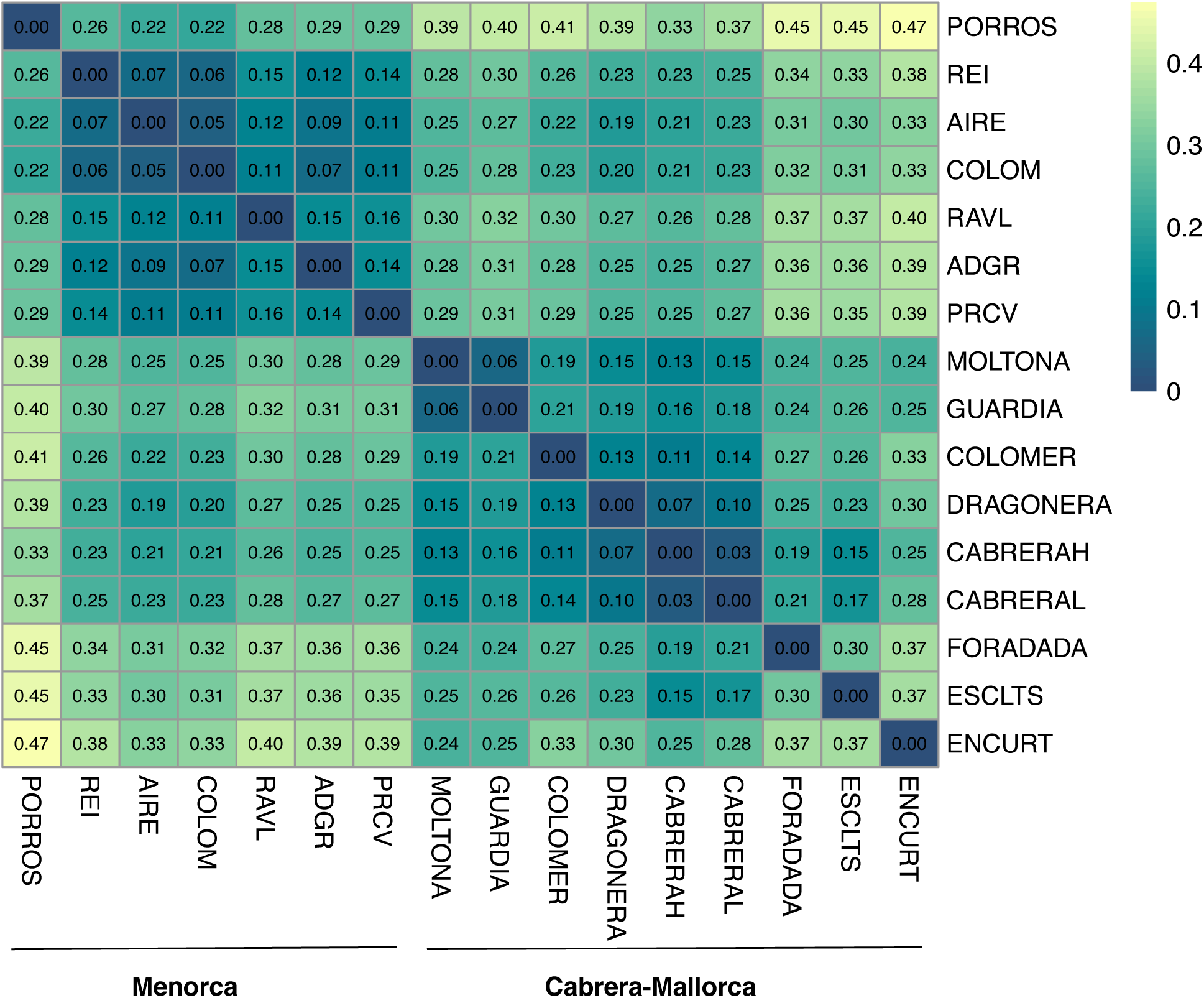
Heatmap of pairwise genetic differentiation (Fst) between the 16 populations of *P. lilfordi* (COMBINED dataset, 2876 SNPs). All Fst estimates were significant (p< 0.01 after permutations).

The most discriminated population was the small islet of en Curt (Mallorca), showing an average Fst of 0.338 with respect to all other islands (minimum value with Moltona, Fst =0.24, Figure 4). The least differentiated populations were Aire and Colom (0.047, Menorca), Na Guardis and Na Moltona (0.064, Mallorca), and the two Cabrera localities (0.030). In this latter case, Fst value <0.05 supports the existence of a unique panmictic population within the major Cabrera Island.

Within major archipelagos, no significant correlation was found between genetic (Fst) and geographic distances (p>0.1), excluding a scenario of isolation-by-distance.

Population genetic clustering with DAPC (Figure 5A) was consistent with Fst-based distances (Figure 4) and the geographic distribution of these populations (Figure 1). Membership probabilities according to the retained discriminant function (64 PCs) correctly assigned all individuals to their source population/islet, excluding recent translocations or mislabeling. According to the first Discriminant Function (DF) (35% of variance), three major genetic clusters were identified (Figure 5A): I) the group of Menorca islands, highly homogeneous and with poor internal discrimination, II) the group of Cabrera and Mallorca islands, characterized by a stronger genetic differentiation among populations, and III) the islet of en Curt, forming an independent lineage. DAPC on the subset of Menorcan populations only (30 PCs retained) slightly increased the population resolution, highlighting a clear divergence of the islet of Porros (Figure 5A, top left panel).

**Figure 5:**
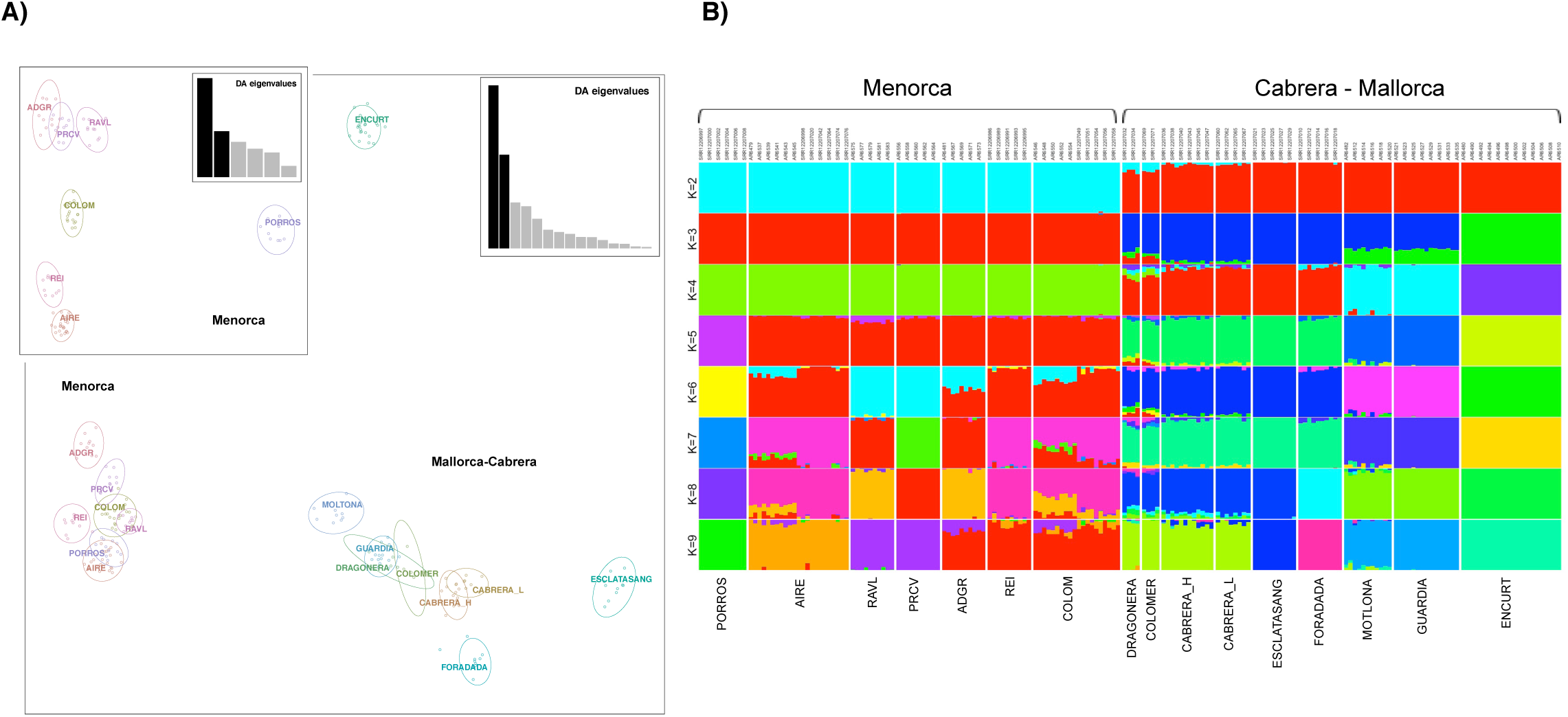
Population genetic structure based on 2876 SNPs (COMBINED dataset, after LD). A) DAPC retaining the first 64 PCs (65% of the explained variance). B) Unsupervised clustering by ADMIXTURE depicting K =2 to K=9 (best value) clusters.

ADMIXTURE analyses were used to further explore population structure, as well as shared ancestry (Figure 5B). Genetic clustering clearly evidenced a major discrimination between Menorca and Cabrera/Mallorca populations (K=2), followed by progressive substructuring within Cabrera/Mallorca (from K=3), and ultimately within Menorca (from K=5). The best-supported number of present genetic clusters is 9 (CV=0.386), corresponding to four genetic clusters in Menorca and five in Mallorca/Cabrera, in line with the major structure recovered by DAPC analysis (Figure 5A). The proportion of shared ancestry among populations varies from zero (independent lineages of the islets of en Curt, Porros, Foradada and Esclatasang) up to 40% (the highly admixed population of Colom). Cabrera island and distant islands from North-West of Mallorca (Dragonera and Colomer) formed a unique genetic cluster, with a small proportion of shared ancestries also with the small islets of Foradada and Esclatasang (North and South of Cabrera, respectively) (see map in Figure 1). According to the optimal current grouping (K=9), no shared ancestry was detected between the two major archipelagos of Menorca and Mallorca/Cabrera, indicating a strong genetic differentiation. However, lower substructuring, from K=2 to K=9 retrieved a clear signature of shared ancestry between the Northern Mallorca populations of Dragonera and Colomer with the Menorcan populations of Colom, Rei and Aire.

### 3.3. Phylogeographic scenario

Prior to phylogenetic reconstruction, we assessed past migration events among populations using allele frequencies on Treemix and OptM analysis to support the best number of edges. OptM analysis indicated a single migration edge explaining 98.9% of the variance. The event with the greatest weight occurred from Colomer into Cabrera-Escatlasang (mean migration weight: 0.454032, mean p-value: 3.22478e-11), followed by a migration event from Dragonera into Escatlasang-Cabrera (mean migration weight: 0.380029, mean p-value: 6.71e-05) (Figure S1).

A midpoint rooted ML tree was then built using consensus sequences for each population on the concatenated SNP dataset (Figure 6). The phylogenetic tree supports the major grouping obtained by the ADMIXTURE, Fst and DAPC analyses, largely in line with the population geographic distribution (Figure 1). Menorcan and Cabrera/Mallorca populations formed clearly distinct phylogenetic clades (bootstrap, bs =100%). Within the Mallorca/Cabrera clade, further substructuring indicated three well differentiated clades: i) the southern Mallorca islets of en Curt, Moltona and Guardia (bs= 99%)), ii) Cabrera islets of Foradada and Esclatasang (bs=76%), with Cabrera as a basal lineage, and iii) the two northern Mallorcan islands of Colomer and Dragonera (bs=80%) (Figure 1 and Figure 6). Within the Menorca clade, substantial support is given only to the grouping of Porros de Cavalleria (PRCV) and Revells (RAVL) (bs=73%), while the small islet of Porros de Fornells (PORROS) represents an independent lineage, highly differentiated from to the rest of Menorcan populations (bs=89%). The remaining Menorcan populations were poorly resolved, consistently with their lower Fst values (Figure 4) and higher admixture levels (Figure 5B).

**Figure 6:**
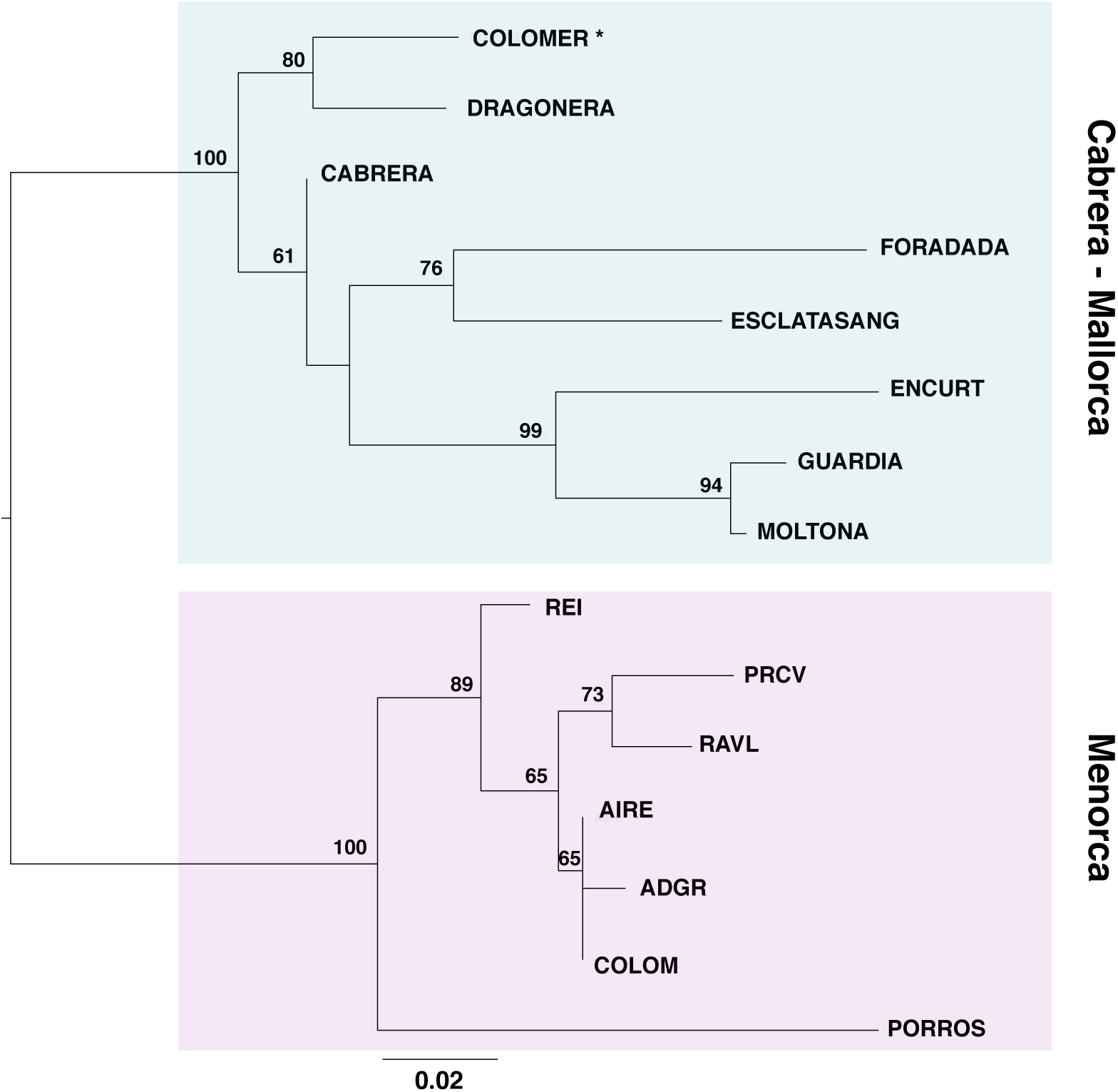
Midpoint-rooted ML tree based on consensus sequences per population after SNP concatenation (2876 SNPs, only variable sites, model GTR+ASC, 10,000 bootstrap replicates). Node numbers indicate bootstrap support. The asterisk indicates the most probable ancestral population according to IQ-TREE 2 root analysis.

Tree rooting using non-reversible models on IQ-tree consistently identified the population of Colomer (Mallorca) as the most likely ancestral lineage. The island showed the highest rootstrap value (86,13%, after 10K bootstrapping on consensus sequences, all sites; values <2% for all other branches) (Figure S2). This result was robust to the use of consensus sequences with variant sites only and to subsampling to one random specimen per population, all consistently recovering the highest rootstrap value for Colomer (71.35 and 69.54, respectively). Root testing provided the maximum p-value of the approximately-unbiased (AU) test for the branch leading to Colomer (p=0.85). All branches within the Menorca clade showed a p<0.05, significantly excluding Menorca as a potential source of the ancestral population. Rootstrap values and root testing considering only the archipelago of Menorca did not provide any clear root support within Menorca.

### 3.4. Outlier identification

We explored potential selective pressures driving population genomic differentiation by identification of outlier SNPs. Intersection of outputs from two independent methods (PCAdapt and Bayescan) provided a total of 18 candidate outliers for the COMBINED dataset, 222 for the GBS and 190 for the RADSeq dataset (n total =446, q<0.01 and *F*_ST_>0.8, see Table S4 for a complete list).

Pattern of distribution of outliers per chromosome was highly concordant among the three datasets, with most outliers falling within the sexual chromosome Z (>16% for all datasets), followed by a large representation on chromosome 2 (>9%) and 15 (12 and 8%, respectively) (Figure 7A). Outliers in chromosomes 3 and 4 were detected only by RADSeq.

**Figure 7:**
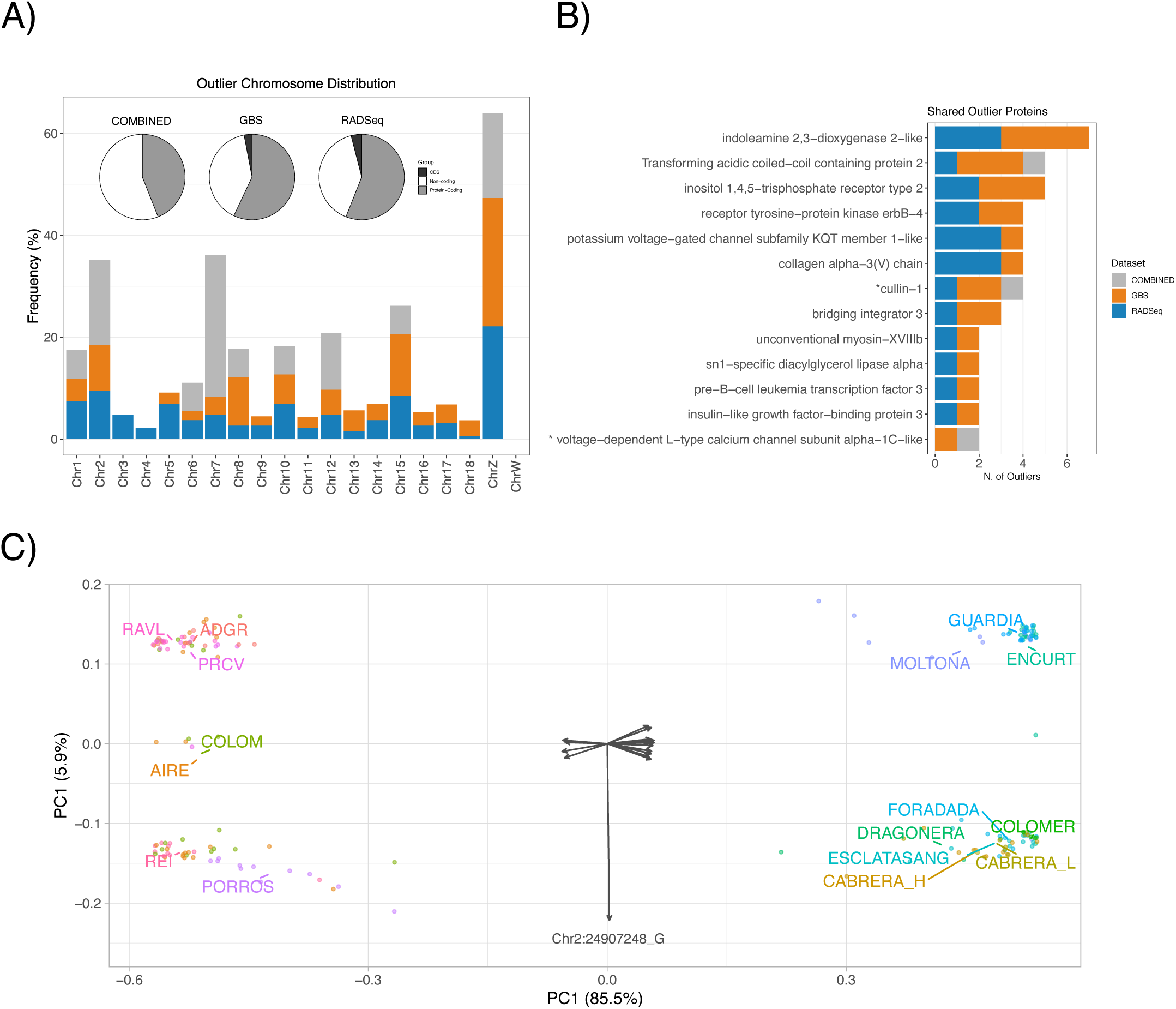
Candidate SNP outliers identified for GBS (n= 222), RADSeq (n=190), and COMBINED (n=18). A) Outliers chromosome distribution (barplot) and proportion per gene category (coding, CDS and non-coding regions) (pie chart). B) Proteins with outliers detected by multiple datasets and corresponding number of outliers. The asterisk indicates proteins with shared SNP outliers among datasets. C) Distance biplot of a Principal Component Analysis (PCA) based on the outlier genotype distance matrix for the COMBINED dataset (18 SNPs). Circles are specimens plot to optimally reproduce their Euclidean distances; arrows are unscaled SNPs scores showing their contribution to each principal component.

About 60% of the outliers fell within protein-coding genes (44% for the COMBINED) corresponding to a total of 182 proteins (Figure 7A, pie charts). Only a minor proportion fell within CDS (n=14 outliers, corresponding to 11 annotated proteins). Five of these SNPs caused a nonsynonymous substitution (ALT) with respect to the reference genome (REF) and were detected in the glypican-3, B-cell lymphoma 6 protein, the cytochrome P450 2C19-like, the pore complex Nup214 genes and an uncharacterized protein (see details in Table S4).

Functional enrichment analysis of annotated proteins with outliers indicated a comparable profile for GBS and RADSeq, showing a significant overrepresentation of binding proteins and catalytic activity (MF category), and metabolic process, growth, and cellular homeostasis (BP category) (BH, p<0.001, Figure S3).

Of the overall 182 proteins with outliers, a small subset (n=13) was consistently retrieved by more than one dataset and most (n=9) were target of multiple outliers per dataset (up to seven) (Figure 7B). Three of these proteins were ion channels: the calcium channels inositol 1,4,5-trisphosphate receptor type 2 and the voltage-dependent L-type calcium channel subunit alpha-1C-like, and the potassium voltage-gated channel subfamily KQT member 1-like. Two proteins presented an outlier position identified by more than one dataset: the cullin-1 protein (SNP Position POS:937545) detected by all three datasets, and the voltage-dependent L-type calcium channel subunit alpha-1C-like (POS:55226525) detected by both GBS and COMBINED (Table S4).

### 3.5. Major Archipelagos-segregating SNPs

Finally, we explored environmental and geographic variables driving outlier distribution. PERMANOVA analyses indicated that Major Archipelago (Menorca and Mallorca/Cabrera, Table 1) was the only explanatory variable for all three datasets (p<0.001, 1000 permutations), with all individual outliers, except one, significantly contributing to geographic between these two major phylogenetic lineages (Figure 6) (binomial test p<0.01). PCA based on outlier genotype matrix clearly evidenced this major clustering (85.5% of variance explained for the COMBINED dataset, Figure 7C, see also Figure S4 for individual outlier genotype clustering). Only one SNP departed from this trend (SNP Chr2_24907248), recovering a clustering of South Mallorca with a subset of the Menorcan islands (Figure 7C and Figure S4). The SNP falls within the sidekick2 protein (Table S4).

A high percentage of outliers had allelic variants that perfectly segregated between major archipelagos, i.e., had null frequency in one archipelago while being present in the other (n total = 154; 11% for COMBINED, 42% for GBS and 31% for RADSeq) (Table S5). These fixed allelic differences can be considered as specific markers of the two major phylogenetic clades (Figure 6).

## 4. Discussion

### 4.1. Genetic diversity of *P. lilfordi*: evolution in islands and population conservation

A fundamental goal in population ecology and evolution is to understand the processes that maintain genetic diversity, and those that drive intraspecific/interpopulation divergence across geographic space and time (Avise, 2000). In small insular populations, free of predators and with no immigrants, genetic diversity is primarily driven by genetic drift (due to environmental and demographic stochasticity, both particularly important in tiny islets) and density-dependent selection due to competition for the limited local resources (Hoffmann et al., 2021; Hunt et al., 2022; Travis et al., 2023). Founder effect and high inbreeding levels are also expected to further reduce the population genetic variance and accelerate the process of divergence from the original source population (Keller and Waller, 2002). Each *P. lilfordi* island clearly hosted a distinct population (all Fst distances were significant), with no contemporary gene flow (no recent translocations detected). Few small islets showed a remarkably high differentiation from all other islets (i.e. Porros and en Curt, Fst >0.2 for all pairs, see Figure 4), indicating either a strong founder effect and/or intense local drift and selection due to their reduced island size (Sendell-Price et al., 2021). Despite being effectively closed, *P. lilfordi* populations harbored relatively high genetic diversity (Ho>0.09 and Pi> 0.1 for all populations), even higher than other continental and insular species of *Podarcis* (Sabolić and Štambuk, 2021). Diversity was overall largely comparable among all 16 populations studied and across their range of distribution (Figure 1 and Figure 3); nonetheless, association analyses supported the expected pattern of decreasing genetic diversity with decreasing island size (Furlan et al., 2012). Smaller islands also presented the highest number of private alleles, indicative of an ongoing process of differentiation. Mallorca/Cabrera archipelago typically hosted a higher genetic diversity than Menorca (Figure 3), with the islands of Cabrera, Guardia and Moltona (South of Mallorca) currently harboring the largest genetic diversity present in this species.

As previously reported by Bassitta *et al*. (2021), we also observed low levels of inbreeding for most populations (Fis< 0.08), even for tiny islets such as Esclatasang, Foradada, Porros, Revells and en Curt (all with Area< 0.7 ha, Fis<0.5, Table 1 and Figure 3). While small populations are expected to show a reduced heterozygosity (Keller and Waller, 2002), Fis values in *P. lilfordi* indicated a slight trend of positive increase with increasing islet surface, with the highest estimate found for the island of Moltona (Fis =0.16). A small inbreeding coefficient, negatively associated with island size, was also found in a previous study based on microsatellites from the three neighboring populations of Moltona, Na Guardia and en Curt (Rotger et al., 2021). A potential explanation is the existence of a population spatial substructure within larger islands, which would cause a non-random mating of individuals effectively reducing the observed heterozygosity. Nonetheless, we invite caution in interpretation of these results as SNP-based estimation of the inbreeding depression in wild populations is notoriously challenging (Schmidt et al., 2021) and would require pedigree-studies for a reliable quantification (Kardos et al., 2016).

Both the relatively high genetic diversity and low Fis values observed in *P. lilfordi* would suggest the existence of potential mechanisms for buffering inbreeding depression in these lizard populations, increasing their chances of persistence. In line with this, recent studies support the ability of insular lizards to counteract genetic depletion even in presence of a strong founder effect (see Sherpa et al., 2023).

### 4.2. A phylogeographic scenario of *P. lilfordi* colonization of the Balearic Islands

*P. lilfordi* genetic diversity was largely structured according to the main geographic distribution, with all analyses supporting the existence of two major genetic lineages separating the large archipelagos of Menorca from Mallorca/Cabrera (Figure 5 and 6). The tree topology was largely consistent with the ones previously reported based on mtDNA and SNPs data (Bassitta et al., 2021; Brown et al., 2008; Pérez-Cembranos et al., 2020; Terrasa et al., 2009). While Mallorca/Cabrera lineage is further sub-structured in four well-supported clades (bs>80%, Figure 6), phylogenetic relationships among Menorcan populations remain largely unresolved, as also indicated by their low degree of differentiation (Figure 4, Fst values < 0.1). Menorcan islands were once largely panmictic (Brown et al., 2008; Terrasa et al., 2009) and a signature of past admixture can still be observed among most populations of this archipelago (Figure 5B), with the exclusion of the small islet of Porros, now representing an independent lineage (Figure 6).

Admixture analysis (K=2, Figure 5B), root inference by IQ-tree (Figure S2) and root testing all strongly suggest that the origin of the species occurred in the main island of Mallorca, with most ancestral populations in our dataset being identified in the North of Mallorca. The highest rootstrap support (86.13%) specifically points to the small island of Colomer as the population retaining nowadays most ancestral polymorphisms, in line with previous proposals (Bassitta et al., 2021; Terrasa et al., 2009). This root placement is consistent with the geographic history of both Dragonera and Colomer islands which were once connected to the Serra of Tramuntana, a mountain chain running up North of Mallorca, before their detachment from the main Mallorca Island about 2.3 Mya (Brown et al., 2008; Terrasa et al., 2009). The Serra of Tramuntana is considered to be the last refuge of *P. lilfordi* (Bailón, 2004) before its extinction in the main Mallorca Island, as testified by fossil records dating around 2000 years old found in the Muleta cave, in the central part of the Tramuntana (Alcover, 2000; Kotsakis, 1981). The Colomer islet is characterized by high lizard density (Pérez-Mellado et al., 2008) and limited accessibility (due to high altitude and coastal shape). Although introduction of lizards from other islands cannot be ruled out, the islet has reduced chances of external gene flow, compatible with the retention of ancestral polymorphisms (Bassitta et al., 2021; Terrasa et al., 2009).

The following process of colonization within the archipelago of Mallorca/Cabrera did not follow an isolation by distance model (Mantel test, p>0.1). According to admixture, phylogenetic analyses and Treemix reconstructions of past migration events, North of Mallorca was then the source of colonization of main Cabrera Island, with which they form a unique genetic clade (see Admixture results, Figure 5B). This closed genetic relationship was also reported by Bassitta et al., 2021, although it partly differs from mtDNA results (Terrasa et al., 2009).

Following this scenario, species expansion within the archipelago of Cabrera proceeded with the colonization of small surrounding islets (Esclatasang and Foradada), as supported by the observed coancestry with Cabrera Island (see Figure 5B). This suggests a founder effect in these small islands, followed by independent evolution leading to the nowadays well-differentiated genetic clade of Esclatasang and Foradada (Figure 5 and 6). Similarly, admixture and ML tree analyses indicated that islets on North of Cabrera could have been a source of colonization for Southern Mallorca islets, specifically the islet of Moltona, with which they still share a small proportion of coancestry (Figure 5B and 6), while the geographically closed sister population of Na Guardis has currently lost all signature of coancestry with Northern Cabrera (Figure 5B). The observed clustering of Cabrera with South Mallorca islets confirms previous haplotype grouping based on mtDNA data, which also suggested a directionality of gene flow from Cabrera to South Mallorca (Network III; Terrasa et al., 2009). Moltona would have then seeded the small islet of en Curt, with which they form a unique clade (bs=99%, Figure 6), as evidenced by their shared coancestry (Figure 5B). The tiny islet of en Curt (0,44 ha) would have followed a process of accelerated differentiation presumably driven by a founder effect, extensive genetic drift, and intense density-dependent selection with more than 1500 ind/ha (Ruiz De Infante Anton et al., 2013). As an alternative scenario, the three islets on the South of Mallorca directly derived from the ancestral mainland Mallorca population, now extinct, a scenario that we presently cannot exclude.

According to bathymetric, geological and genetic data subsequent colonization of Menorca occurred around ∼ 2.8 Mya (Brown et al., 2008; Terrasa et al., 2009). We were not able to assess the root within this archipelago due to high levels of admixture. Menorcan populations were highly panmictic during a large period of time and the colonization and subsequent isolation of most Menorcan islands was gradual (Pretus et al., 2004), following a vicariance type of colonization, with an asymmetric flow from south to north of the distribution of the species (Terrasa et al., 2009).

### 4.4. Signature of genomic diversification of *P. lilfordi* populations

Local adaptation in effectively closed populations is a trade-off between genetic drift and selective pressures (Savolainen et al., 2013). The two processes are intrinsically associated with the island surface and its exposure to open sea, both affecting the impact of environmental stochasticity (stronger in smaller islets) and resource availability (reduced in smaller islands). Particularly, the vegetation cover, highly correlated to the island size, provides a strong cascade effect on lizard resource availability, including pollen and fruits production, insect visitor frequency and diversity, and even seabirds presence, all potential sources of dietary items (Pérez-Mellado et al., 2008; Ruiz De Infante Anton et al., 2013; Salvador, 2009; Santamaría et al., 2020). Along with stochasticity, this gives rise to a heterogeneous landscape across islands, which is expected to drive independent processes of lizard phenotypic diversification.

We explored the signature of genomic diversification that might underpin this lizard phenotypic diversity through outlier analysis. Outlier distribution showed a clearly skewed chromosome representation, with most outliers falling within the sexual chromosome Z, consistently for all datasets analyzed (Figure 7A). Recent studies in birds and lizards showed that closely related species often present high differentiation on the Z chromosome (Kulikova et al., 2022; Rovatsos et al., 2019), a pattern that is typically explained as faster Z evolution and lower recombination rates (Irwin, 2018; Lima, 2014; Mank et al., 2010; Wright et al., 2015). The existence of potential "islands of differentiation" within the sexual chromosomes has also been hypothesized (Lavretsky et al., 2019). At present, detection of such “islands” in *P. lilfordi* Z chromosome would require a higher sequencing coverage than the one currently provided by GBS and RADseq (i.e., genome resequencing).

Most outliers fell within protein-coding genes, although the large majority were intronic (Figure 7A). Substantial evidence supports the notion that introns have a crucial and evolutionarily conserved function in controlling gene expression in eukaryotes (Kumari et al., 2022; Rose, 2019), providing a particularly rapid mechanism for increasing variation in proteome variance by production of a diverse array of alternative splicing variants (AS) (Reixachs-Solé and Eyras, 2022; Wang et al., 2015). Moreover, most genes with outliers were associated with protein binding and catalytic activities and were involved in metabolic and growth processes (Figure S3), critical molecular functions for gene expression regulation and phenotypic diversification (Van Nostrand et al., 2020). In small, isolated populations, extensive modulation of gene expression conferring phenotypic plasticity could represent a major mechanism to counteract the loss of genetic diversity (Fulgione et al., 2023; Sherpa et al., 2023), a hypothesis that needs to be validated by whole transcriptome and epigenome data (Chapelle and Silvestre, 2022; Fulgione et al., 2023).

We found a subset of protein-coding genes targets of multiple outliers according to both independent sequencing methods, GBS and RADSeq (Figure 7B), which might represent interesting candidates for further exploration of population genomic diversification in *P. lilfordi*. Of these proteins, few represented potassium and calcium channels, which could hint to an important regulatory role of osmotic pressure in lizards (Dantzler and Bradshaw, 2008). We also highlight the cullin-1 protein, with a shared outlier SNP detected by all datasets. The protein is known to be involved in ubiquitination and subsequent proteasomal degradation of target proteins (Duan et al., 2011; Gao et al., 2011; Scott et al., 2016). Recent evidence indicates a critical role of this protein in modulation of the transcription factor c-MYC, a major regulator of gene expression and cell proliferation (Sweeney et al., 2020). While there are no specific studies on cullin-1 protein in lizards, c-MYC was linked to the cellular regenerative response after tail amputation (Alibardi, 2017). The gradual loss of tail autotomy ability in insular lizards is a hallmark of their reduced antipredator response following insular adaptation (part of the island syndrome) (Cooper et al., 2004; Pafilis et al., 2009; Perez-Mellado, 1997).

Exploration of the major variables driving outlier genotypes primarily recovered a signature of past geographic separation between Menorca and Cabrera/Mallorca, with a large proportion of loci presenting fixed allelic differences between archipelagos (Figure 7C). These SNPs are putatively derived from a past founder effect that occurred during initial colonization of the Menorca archipelago (Brown et al., 2008; Terrasa et al., 2009) or the result of unclear selective forces, including potential selective sweeps (Brown et al., 2023; Campagna et al., 2022; Stephan, 2019). In all cases, they support no recent secondary contact between Archipelagos, in line with previous mtDNA-based phylogeographic reconstructions (Terrasa et al., 2009). We note that given the major confounding effect of archipelago, the significance/impact of other environmental variables in driving genome diversification cannot be reliably assessed with the current sampling design (Bassitta et al. 2021). Additional sampling, along with phenotypic data are required to clarify the putative adaptive role of these genomic changes.

## Conclusions

The Balearic lizard *P. lilfordi* has recently entered the genomic era, offering a new interesting insular system to explore trends in vertebrate genome evolution. Our study presents a first use of the *P. lilfordi* genome to reliably mapped extensive SNP data. Comparative analysis using independent methods and datasets provided robustness to our findings. We found support for the origin of this species colonization in the Mallorca Island, confirming previous proposals and providing a framework for future studies of the evolutionary trajectories in genome diversification for this insular species. The substantial genetic diversity observed in these effectively closed populations suggest that they have potential mechanisms to partially counteract genetic drift and inbreeding depression that are worth further investigation by genome resequencing and inclusion of additional populations. Furthermore, integration of population demographic data (ongoing) and collection of detailed phenotypic data, still scarce for this species, will be critical to evaluate the genomic plasticity and ability to persist of these insular populations.

## Supplementary data

**Figure S1:**
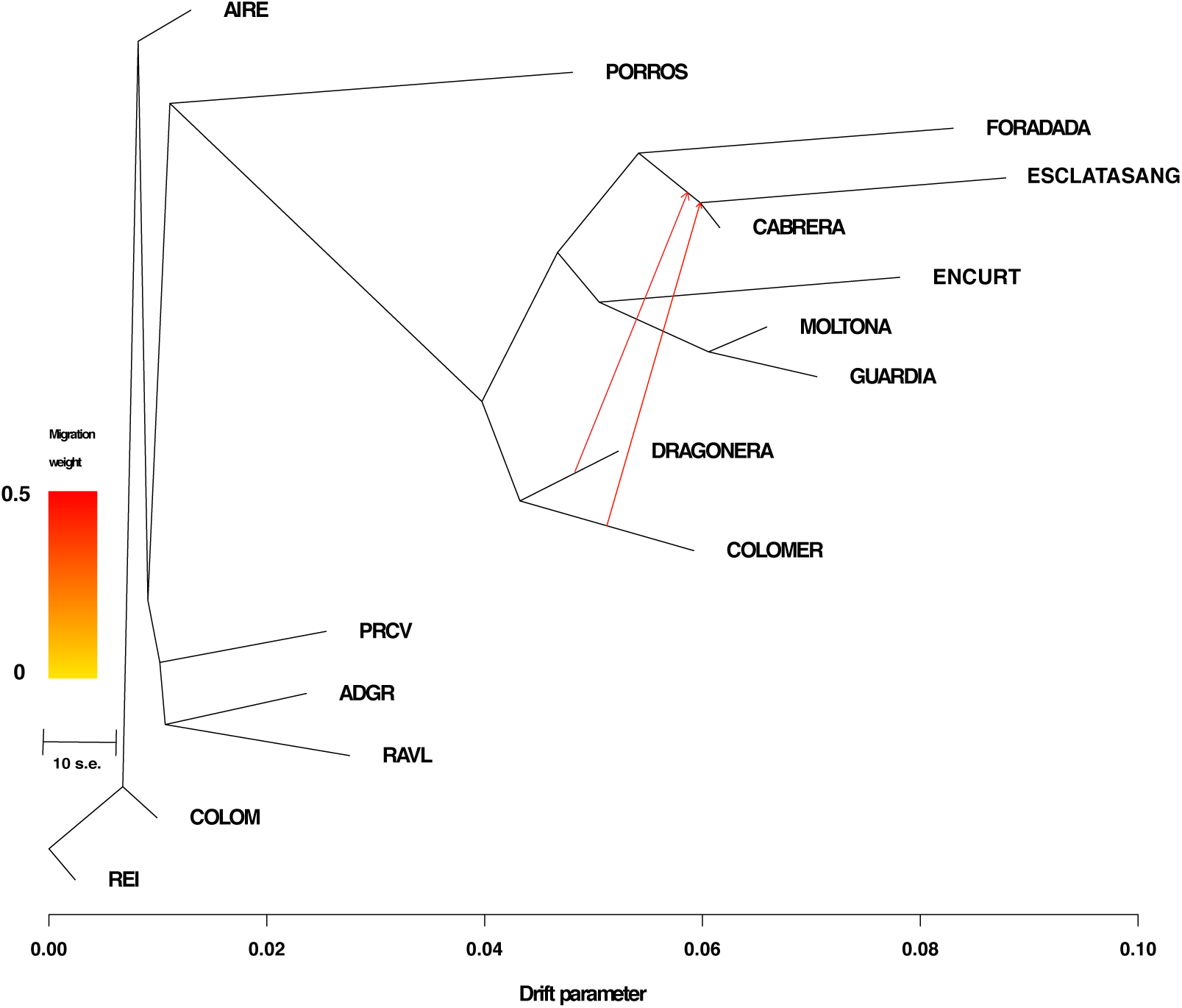
The ML-graph inferred by TreeMix with 16 migration events based on the COMBINED dataset. The arrow indicates the two significant migration events with the highest migration weight. Drift parameter is shown on the x-axis.

**Figure S2:**
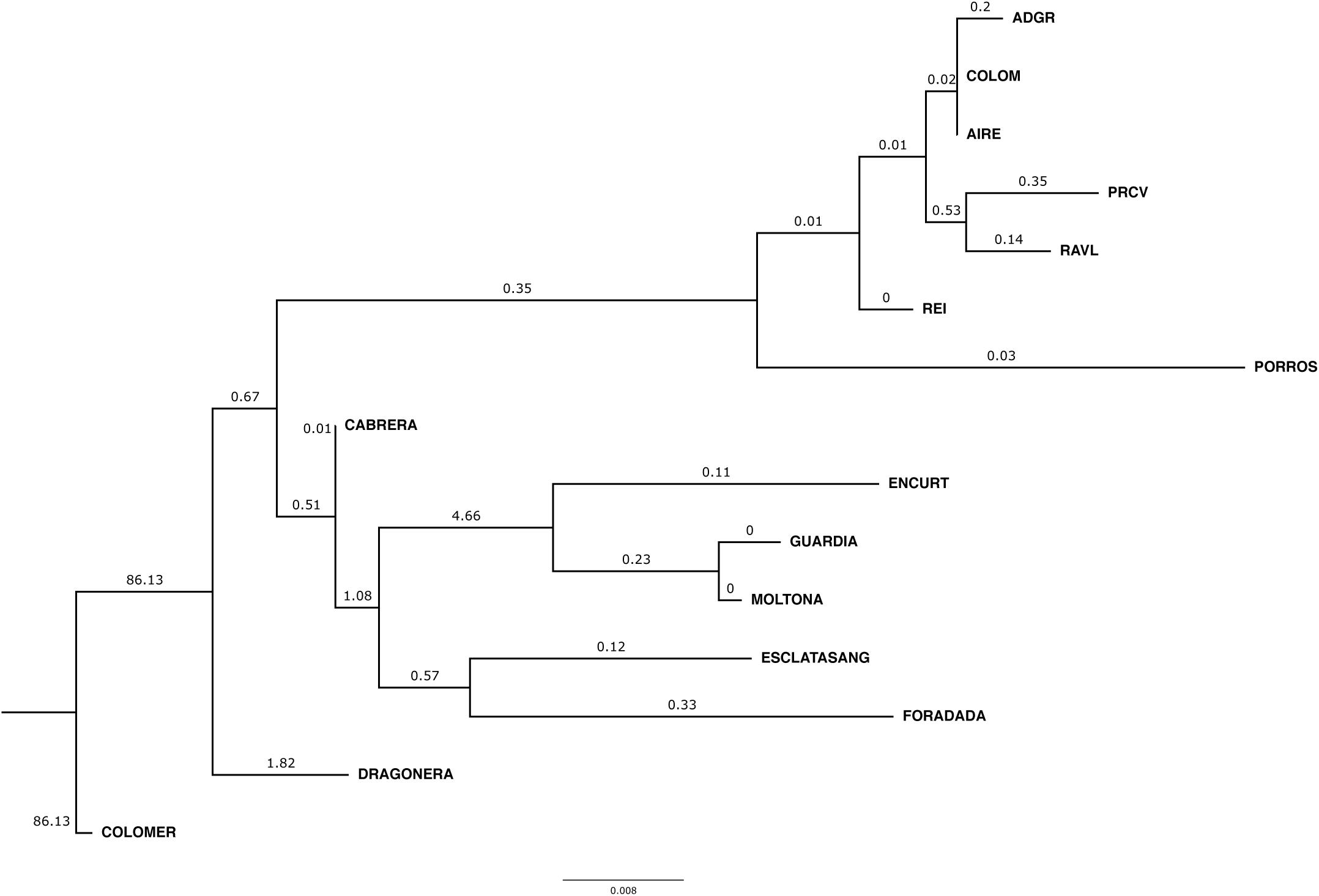
ML tree with *rootstrap* support values for root inference. Values were estimated with IQ-TREE 2 on consensus sequences per population (2867 bp), using a non-reversible model (model 12.12, all sites, 10000 bootstrap replicates).

**Figure S3.**
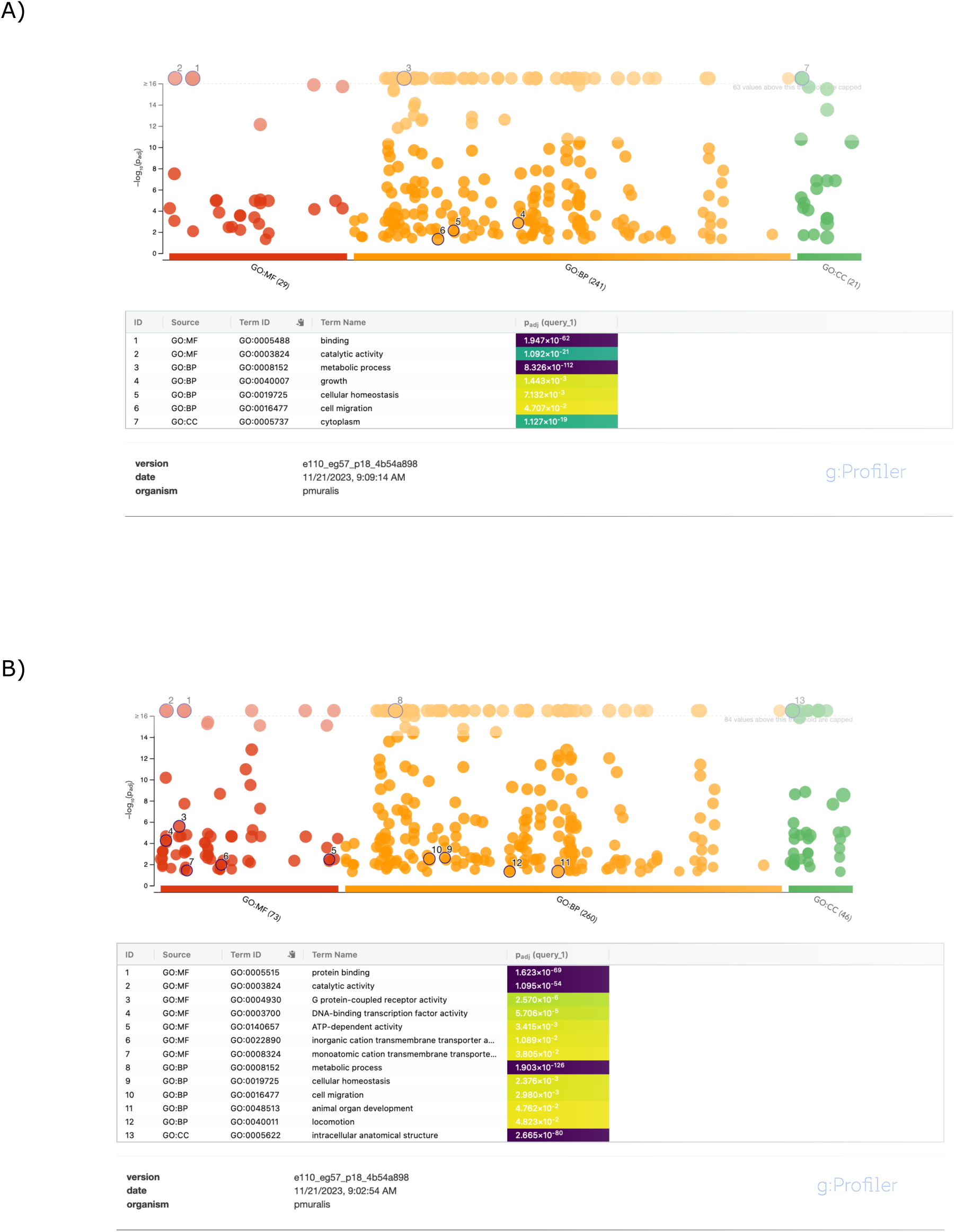
Functional enrichment of GOs for proteins with outliers, according to A) GBS and B) RADSeq dataset.

**Figure S4.**
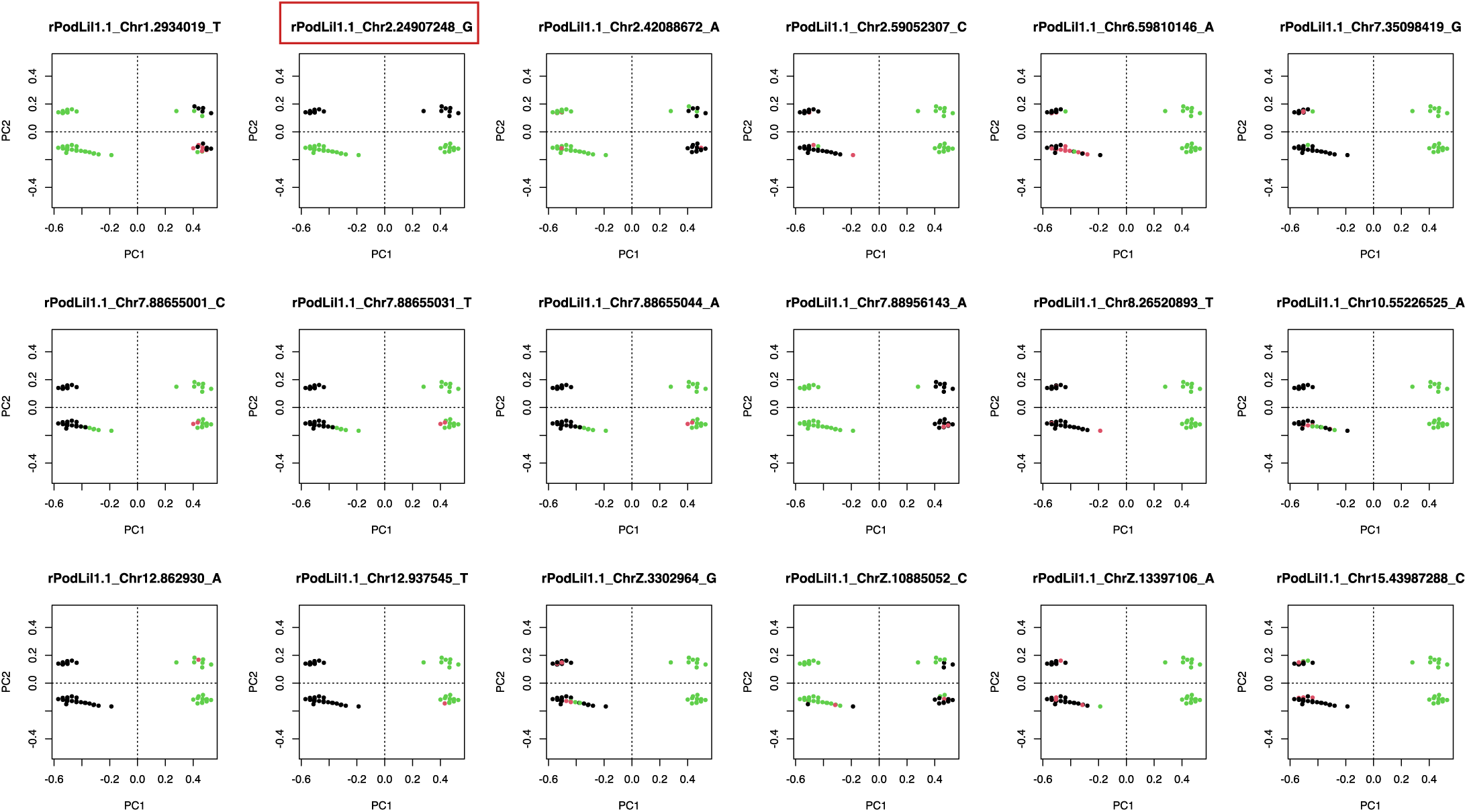
PCA of the 18 outliers detected in the COMBINED dataset. All but one indicated a clear clustering by archipelago. Black dots correspond to the most frequent allele, while Red dots indicate the heterozygotic genotype.

**Table S1:** Individual sample metadata.

**Table S2:** GBS read summary.

**Table S3:** Genetic diversity estimates for individual datasets (GBS and RADSeq).

**Table S4:** List of outliers identified by both Bayescan and PCADAPT (Fst>0.8, q<0.01,) for the three datasets: COMBINED (N=18), GBS (222) and RADSEQ (190) and corresponding annotation.

**Table S5:** Archipelago-segregating SNPs (fixed alleles).

## Data availability

The raw data with the individual sequences are available at the Sequence Read Archive (SRA) (BioProject ID: PRJNA1070579).

## Acknowledgments

This study was supported by the CAIB - Government of the Balearic Islands (project PRD2018/25 to G.T.), the Institut d’Estudis Catalans under the Catalan Biogenome Project initiative (PRO2020-S02 to L.B.), the MCIN/AEI/10.13039/501100011033/FEDER, UE (PID2022-141578NB-C22 to L.B.) and the Doctoral Scholarship, Colombian Ministry of Science, Technology and Innovation (MINCIENCIAS885/2020 to K.O)

